# Molecular mechanisms of catalytic inhibition for active site mutations in glucose-6-phosphatase catalytic subunit 1 linked to glycogen storage disease

**DOI:** 10.1101/2023.03.13.532485

**Authors:** Matt Sinclair, Richard A Stein, Jonathan H Sheehan, Emily M Hawes, Richard M O’Brien, Emad Tajkhorshid, Derek P Claxton

## Abstract

Mediating the terminal reaction of gluconeogenesis and glycogenolysis, the integral membrane protein G6PC1 regulates hepatic glucose production by catalyzing hydrolysis of glucose-6-phosphate (G6P) within the lumen of the endoplasmic reticulum. Consistent with its vital contribution to glucose homeostasis, inactivating mutations in G6PC1 cause glycogen storage disease (GSD) type 1a characterized by hepatomegaly and severe hypoglycemia. Despite its physiological importance, the structural basis of G6P binding to G6PC1 and the molecular disruptions induced by missense mutations within the active site that give rise to GSD type 1a are unknown. Exploiting a computational model of G6PC1 derived from the groundbreaking structure prediction algorithm AlphaFold2 (AF2), we combine molecular dynamics (MD) simulations and computational predictions of thermodynamic stability with a robust *in vitro* screening platform to define the atomic interactions governing G6P binding as well as explore the energetic perturbations imposed by disease-linked variants. We identify a collection of side chains, including conserved residues from the signature phosphatidic acid phosphatase motif, that contribute to a hydrogen bonding and van der Waals network stabilizing G6P in the active site. Introduction of GSD type 1a mutations into the G6PC1 sequence elicits changes in G6P binding energy, thermostability and structural properties, suggesting multiple pathways of catalytic impairment. Our results, which corroborate the high quality of the AF2 model as a guide for experimental design and to interpret outcomes, not only confirm active site structural organization but also suggest novel mechanistic contributions of catalytic and non-catalytic side chains.

## Introduction

Interconversion of glucose between its free and phosphorylated forms is among the most central reactions in metabolism, controlling its transport between extracellular, cytoplasmic, and luminal compartments, and thereby its storage and homeostasis. For example, glucose-6-phosphatase (G6Pase) couples the uptake of intracellular glucose-6-phosphate (G6P) into the endoplasmic reticulum (ER) lumen (1) with G6P hydrolysis into glucose and inorganic phosphate (Pi) (2, 3). The hydrolysis reaction is catalyzed by one of three membrane-embedded G6Pase catalytic subunit (G6PC) family members that differ in their tissue expression patterns (4). The terminal reaction of gluconeogenesis and glycogenolysis is mediated by G6PC1, which is predominantly expressed in the liver, kidney and small intestines (5–7). Consequently, G6PC1 serves as the gatekeeper for hepatic glucose production that stabilizes blood glucose between meals. By residing at the crux of glucose homeostasis, G6PC1 is a potential therapeutic target for disorders of glucose metabolism. Whereas upregulation of *G6PC1* gene expression contributes to diabetes by increasing fasting blood glucose concentration (8–17), aberrant mutations in G6PC1 that impair activity cause glycogen storage disease (GSD) type 1a (18, 19).

With an approximate incidence of 1 in 100,000 births, GSD type 1a is an autosomal recessive disorder characterized primarily by severe hypoglycemia following a fast (20). Due to inactive G6PC1, the accumulation of intracellular G6P stimulates flux through alternate metabolic pathways including glycogenesis that promotes excessive glycogen storage in the liver. In addition to hepatomegaly, GSD type 1a is also associated with a host of other metabolic complications including hyperuricemia and hyperlipidemia and an enhanced risk of hepatic adenoma and carcinoma (21–23). The disorder is juvenile lethal if left untreated. Although adenoviral gene delivery and lipid-encapsulated mRNA are being explored as clinical treatment options (24–26), therapeutic interventions are limited presently to dietary restrictions supplemented with a drug regimen to treat the disease complications (27). According to the Human Gene Mutation Database, more than 100 pathogenic missense/nonsense mutations scattered throughout the *g6pc1* coding sequence have been identified from GSD type 1a patients (28). However, the structural and mechanistic basis for catalytic dysfunction of these mutations remains to be elucidated.

Despite considerable effort since its discovery in the early 1950s (7), G6PC1 extracted from its native membrane environment has been recalcitrant to *in vitro* investigations due to structural and catalytic instability (29–32), precluding detailed analysis of the structure/function paradigm. In 2021, DeepMind revolutionized the field of structural biology with the introduction of AlphaFold2 (AF2), a deep learning platform that integrates knowledge of protein folds with residue co-evolution encoded within deep multiple sequence alignments to predict remarkably accurate structures of proteins (33). Highlighted in one of two seminal publications (34), the AF2 model of G6PC1 displayed a fold consistent with the type 2 phosphatidic acid phosphatase (PAP2) superfamily that includes membrane-integrated lipid phosphatases and water-soluble haloperoxidases (35) as well as the predicted topology of G6PC1 from biochemical experiments (36, 37). The defining feature of this superfamily is a signature tripartite sequence motif (KX_6_RP---PSGH---SRX_5_HX_3_D/Q) that participates in active site formation (38) and includes positions of GSD type 1a mutations (18). Although the AF2 model recasts G6PC1 and disease-linked mutations in a novel structural context, notably absent from the computational prediction is an understanding of fundamental substrate binding interactions critical to the catalytic mechanism.

In this work, we use molecular dynamics (MD) simulations to explore G6P binding to the G6PC1 AF2 model immersed in a representative ER membrane. Moreover, we predict the molecular consequences to G6P binding energetics and G6PC1 thermodynamics by introducing a select panel of GSD type 1a mutations into the active site *in silico*. Capitalizing on our recently published *in vitro* methods that overcome previous experimental limitations with purified enzyme (39), we complement the computational evaluation of the AF2 model with biochemical and biophysical studies of G6PC1 variants to outline changes in expression, activity, and stability. We find that the AF2-generated model is remarkably stable and binds both α- and β-G6P anomers with a nearly identical pattern of electrostatic and van der Waals interactions within the predicted active site. Perturbation of these interactions through disease-linked mutations not only reveals changes to G6P binding energetics but also to G6PC1 structure and intrinsic stability. The integrated analysis suggests novel side chain contributions to substrate binding and Pi release, and implies that GSD type 1a mutations disrupt catalysis via unique molecular pathways.

## Results

### Structure and stability of G6PC1 in a model lipid bilayer

We chose to exploit mouse G6PC1 as an archetype G6Pase catalytic subunit for the structural and mechanistic questions addressed here. Mouse G6PC1 bears 89% sequence identity with the human homolog and is predicted by AF2 to possess an identical tertiary fold (root mean squared deviation ∼0.125 Å) epitomized by a conserved active site organization. Globally, 65 of the 69 amino acids at positions of naturally occurring GSD type 1a missense/nonsense mutations, including all of those at the active site, are strictly conserved between the two homologs. Moreover, a murine model of GSD type 1a (*g6pc1*^-/-^) recapitulates a similar phenotype as human patients (40, 41). Importantly, detergent-solubilized mouse G6PC1 demonstrates elevated structural and catalytic stability relative to human G6PC1 in biochemical assays (39), supporting an unequivocal interpretation of the *in vitro* studies reported here. Thus, mouse G6PC1, subsequently referred to simply as G6PC1, is an appropriate model for computational and experimental studies.

Because the AF2 modeling pipeline does not account explicitly for the membrane, the G6PC1 model was placed into an approximate ER membrane to assess its relative stability (Fig 1a). G6PC1 was almost entirely buried within the membrane with the active site residing at the membrane-water interface. This model, which was gently relaxed from its initial state during the minimization phase, was then subjected to equilibrium MD simulations, which converged within 100 ns in multiple iterations to arrive at a common structural model suitable for ligand docking and thermodynamic analysis. The aggregate simulation data yielded a final all-atom RMSD of 1.76 Å relative to the original AF2 model (Fig 1b), consistent with a highly stable structure within experimental error (33, 42). Regions of higher internal flexibility are highlighted by root mean squared fluctuation (RMSF) analysis of the simulation dataset (Fig 1c) and correlate well with unstructured loops and regions of lower confidence from the AF2 prediction.

**Figure 1.**
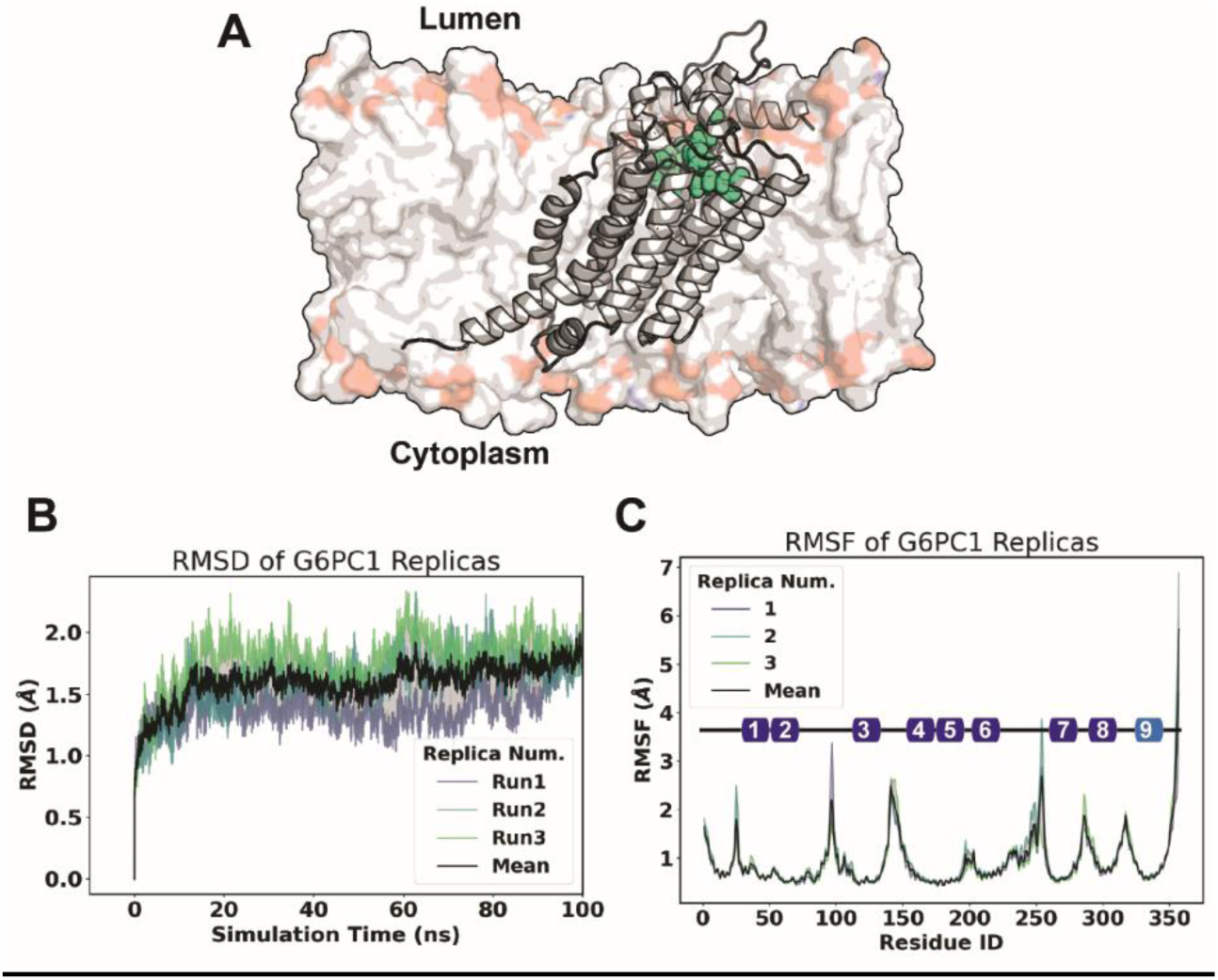
Modeling the AF2 G6PC1 structure in a simulated bilayer. (A) Cartoon rendering of G6PC1 immersed into a representative ER bilayer. The composition is reported in Methods. The active site is demarcated by residues contributing to the phosphatidic acid phosphatase sequence motif (green). (B) Time series of RMSD of G6PC1 over the course of equilibrium simulations. (C) RMSF showcases regions of higher disorder that map to termini and loops. The colored boxes illustrate the topology of the nine transmembrane helices along the primary sequence. The color represents the pLDDT score of the AF2 model for these segments, with the darker blue indicating higher pLDDT that reflects higher accuracy of the model.

### Modeling the G6P-bound active site

We leveraged available crystallographic structures from one vanadium-containing chloroperoxidase (43) and three bacterial PAP2 type lipid phosphatases (44–46) that contain surrogate phosphate molecules (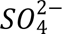, 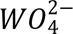, and 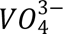) to guide docking of the G6P substrate into the putative G6PC1 active site. When aligned with the AF2 model of G6PC1, these protein structures highlight a strongly conserved active site organization composed of key residues in the phosphatase sequence motif (SI Appendix, Fig. S1a). We mapped the phosphate moiety of both α- and β-G6P anomers, derived from 95 crystal structures, to the AF2 G6PC1 model by targeting the atomic coordinates belonging to the phosphate analogs. Although placement of the phosphate group for each modeled G6P was well defined, a “cloud” of possible conformations was showcased by the glucose moiety likely due to dihedral flexibility (SI Appendix, Fig. S1b, e).

Two conformations representing an α- and a β-G6P anomer were chosen for preliminary docking onto the bilayer-relaxed G6PC1 structure (SI Appendix, Fig. S1c). These two G6P-bound models were subjected to additional 250 ns equilibrium MD simulations performed in triplicate to assess stability. Although repacking of critical active-site residues was observed early in the simulations (SI Appendix, Fig. S1d), RMSD analysis indicated stability and robustness similar to the Apo state (Fig 2a). Both G6P anomers induced marginal structural perturbations relative to the Apo model (< 2.5 Å RMSD). Remarkably, free energy perturbation calculations indicated that binding of the β anomer is more favorable than α-G6P by ∼5 kcal/mol, which may reflect more stabilizing interactions with G6PC1 facilitated by the lower energy configuration of the anomeric carbon (SI Appendix, Fig S2). In some cases, residues lining the active site differed in contact probability between anomers, but these differences were not significant at the 0.05 level for the current dataset (Fig 2b).

**Figure 2.**
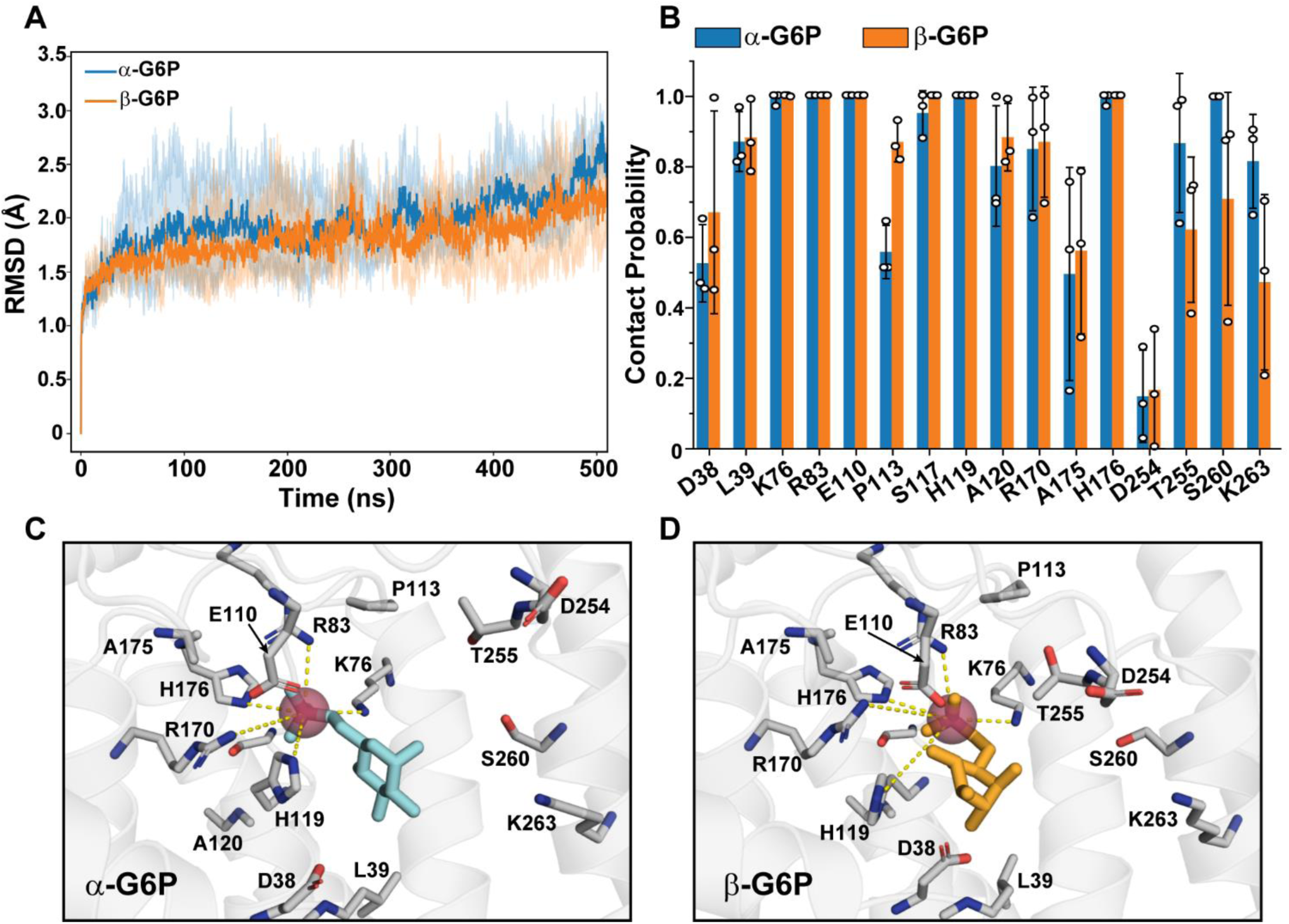
Identification of G6P binding residues and stability of the ligand-bound model. (A) RMSD of substrate-bound simulations for each anomer. The dark traces are the average RMSD over multiple runs (n=3). (B) Per residue contact probability, defined as the proportion of time each residue spends within 4 Å of G6P, was assayed from over 10,000 frames. (C-D) Representative binding modes of α-G6P (C) or β-G6P (D) to G6PC1. The pose chosen was based on the shortest contact distance to side chains of the phosphatase sequence motif.

Fig 2c-d captures a representative binding mode sampled for each G6P anomer over the course of the simulations as determined by a minimal sum of distances between the catalytic K76, R83/170, and H119/176 side chains. For these poses, atomic distances between polarized hydrogens of the side chain functional groups and the phosphate oxygen atoms of both anomers typically spanned < 2-4 Å. Interestingly, the backbone amides of G118 and H119 were in good position to provide hydrogen bonding contacts with phosphate moiety whereas the H119 side chain supported coordination of the glucose ring. Potential longer range (> 4 Å) interactions between the phosphate moiety and backbone amides of S117 and A120 were also observed (SI Appendix, Fig 3). Collectively, the structural convergence of multiple independent simulations as well as the intricate and consistent pattern of interactions with G6P strongly support the predicted active site identity of G6PC1.

### Energetic perturbations arising from clinically relevant mutations

Nine of the 16 residues that make contacts with G6P over the simulation time course are sites of naturally occurring GSD type 1a mutations in G6PC1 (Table 1). Previous *in vitro* characterization of these missense variants in the context of ER microsomes showed that G6P catalysis was reduced or abrogated (19), but the molecular basis of compromised function was unclear. Based on the G6P-bound model alone, mutation of interacting residues may affect substrate binding energy. Alternatively, side chain substitutions may destabilize critical packing interactions. Calculation of changes in Gibbs energy (ΔΔG) within an implicit membrane using the Rosetta macromolecular modeling suite (47–50) predicted that six of these variants (D38V, K76N, R83H, E110K, P113L and T255I) promoted thermodynamic instability greater than 2 kcal/mol in the bilayer-relaxed Apo model (Table 1, SI Appendix, Fig S4). By contrast, H119L and R170Q enhanced stability. A similar predicted pattern was observed using an orthogonal method that calculates ΔΔG from energy-minimized AF2 structure predictions (51), although the absolute values differed (Table 1). The significance of these findings was emphasized by ΔΔG analysis of two uncharacterized missense variants, Q14R and E319K, listed in the Genome Aggregation Database (52) that are found outside the active site on the protein surface and expected to be well tolerated (Table 1). These predicted thermodynamic perturbations suggest that compromising native side chain energetic interactions could disrupt active site stability, potentially contributing to local or global misfolding. We further explored these possibilities by a series of computational and biochemical assays.

**Table 1.** Stability of G6PC1 variants.

| **Variant** | **Clinical Significance<sup>a</sup>** | **$\Delta\Delta G$-Ros (kcal/mol)<sup>b</sup>** | **$\Delta\Delta G$-AF (kcal/mol)<sup>c</sup>** | **T<sub>M</sub> (°C)** |
| --- | --- | --- | --- | --- |
| Q14R | Unknown | +1.0 | +0.6 | 38.8 |
| D38V | Pathogenic | +3.7 | +2.5 | 33.0 |
| K76N | Pathogenic | +2.8 | +1.3 | 36.7 |
| R83H | Pathogenic | +11.7 | +6.3 | N.D. |
| E110K | Likely Pathogenic | +7.0 | +1.4 | N.D. |
| P113L | Pathogenic | +161.0 | +0.6 | N.D. |
| G118S | Pathogenic | -0.7 | +0.4 | 36.5 |
| H119L | Pathogenic | -4.0 | -1.5 | 37.8 |
| R170Q | Pathogenic | -5.5 | -8.8 | 41.7 |
| T255I | Pathogenic | +71.5 | +3.8 | 36.3 |
| E319K | Unknown | +0.8 | +1.9 | N.D. |

<sup>a</sup>ClinVar (82), gnomAD (52) and HGMD (28) genomic databases
<sup>b</sup>Calculated with the bilayer-relaxed Apo model in Rosetta
<sup>c</sup>Calculated from Rosetta energy-minimized AF2 predictions

Residue-specific interaction energies with both G6P anomers were calculated using NAMD (53, 54). The time-series of energy distributions were binned from three independent 250-ns replicates for a total of 54 mutant simulations accumulating 13.5 µs of sampling. The consequences of the *in silico* mutagenesis on G6P binding are shown in Figure 3. Although the absolute number of contact events was reduced across all variants, interaction energies could be categorized into three classes. In one class, the enthalpic component to G6P interactions was largely abolished by non-conservative substitution of key phosphatase residues K76, R83, H119 or R170 incurring ∼20-100 kcal/mol loss of interaction energy in each. According to the model, these positively-charged native side chains are positioned optimally to coordinate the negatively-charged phosphate moiety of G6P (Fig 2). In a second class, the introduction of the D38V, P113L, G118S or T255I mutations was less impactful (though non-negligible) by demonstrating similar energy distributions, suggesting that impaired function of these variants originates from either a distinct mechanism or a combination of factors.

**Figure 3.**
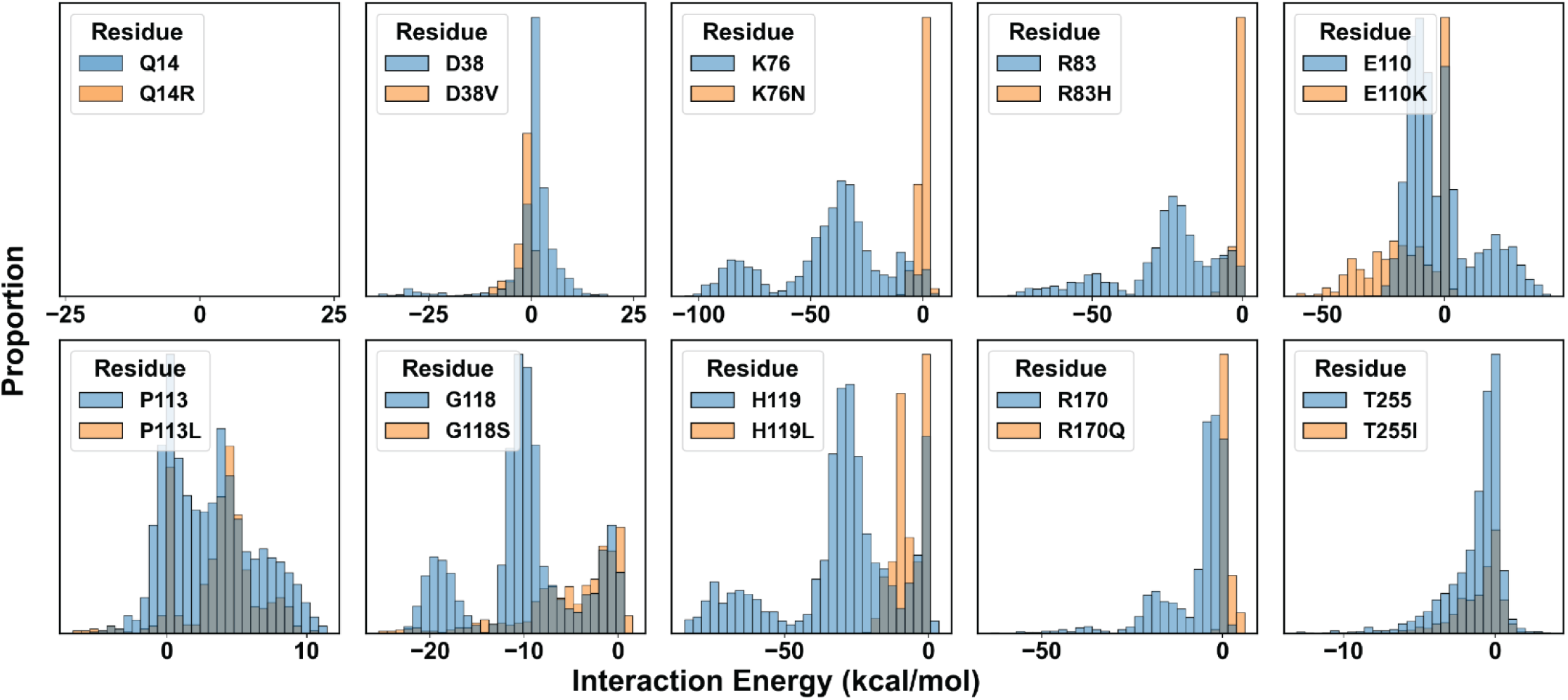
Perturbations of non-bonded interaction energies with G6P in GSD type1a variants. Interaction energy of each residue with G6P illustrates changes in the enthalpy of binding. Data are pooled from both G6P anomers. Energies shown are the summation of van der Waals and electrostatics. Q14R, a negative control, sits outside the active site and does not interact with G6P.

The distinct pattern emerging from E110K established a third class defined by a left shift in the energy histogram indicating greater stabilization of the substrate in the binding pocket. Recently, E110 was postulated to serve as a surrogate substrate of the Apo enzyme by stabilizing a closed conformation of the active site through salt bridges with the nearby R83 and R170 (34). Consistent with this hypothesis, hydrogen bonding was observed between E110 and a polarized hydrogen belonging to the phosphate moiety of G6P of both anomers, shifting the ensemble of salt bridge dynamics (SI Appendix, Fig S5). While this electrostatic interaction with the substrate attenuated salt bridge potential of E110 with R83, interaction of E110 with R170 increased (SI Appendix, Fig S5). Importantly, the MD simulations indicated that K110 is more energetically favorable to G6P binding than E110, which is likely rooted in the increased positivity of the binding pocket and the disrupted salt bridge interactions with the conserved Arg side chains. Thus, these observations imply that while the native Glu (E) participates in product release, the E110K variant favors trapping of G6P in the active site.

### Expression and activity profiles of GSD type 1a variants

To explore the hypotheses derived from our computational analysis, we tested the biological outcomes of these active-site GSD type 1a mutations as well as the Q14R and E319K variants by employing the enzyme expression and activity screens that we developed recently (39). Specifically, WT and mutant G6PC1 were transfected into adherent HEK293SG cells as a fusion with EGFP on the C-terminus to facilitate assay analyses. G6PC1 expression was confirmed via cell epifluorescence (SI Appendix, Fig S6). Relative quantities and homogeneity of G6PC1 fusion protein were determined by EGFP fluorescence detection size exclusion chromatography (FSEC) (55) following whole cell solubilization with lauryl maltose neopentyl glycol (LMNG) detergent. LMNG was identified previously from a detergent screen to support extraction of active and homogeneous G6PC1 (39).

Representative FSEC traces in Figure 4a illustrate the spectrum of chromatographic behaviors observed for the variants. The full complement of traces is shown in SI Appendix Fig S7. In general, expression of disease-linked variants was either unperturbed or greatly reduced relative to WT. Additionally, the elution profiles, which indicate the presence of either polydisperse entities or folded monodisperse protein, reported a variable propensity for aggregation. The R83H and P113L mutations displayed highly compromised G6PC1 expression with elution profiles consistent with a large degree of structural disorder. Although D38V and E110K expression levels were attenuated strongly, the peak shapes suggested that these mutants largely retained the structural properties of the WT. Interestingly, K76N, G118S, H119L, R170Q and T255I behaved similarly to WT, Q14R, and E319K. These features were captured by integration of the elution peak (AUC, area under the curve) from multiple independent trials, translating the FSEC traces into an expression bar graph for each variant relative to WT (Fig 4b). The observed trends were comparable to the whole cell epifluorescence pattern (SI Appendix, Fig S6). Statistical analyses of these data supported the conclusion that expression of Q14R, K76N, G118S, H119L, R170Q, T255I and E319K was unchanged relative to WT (SI Appendix, Table S1).

**Figure 4.**
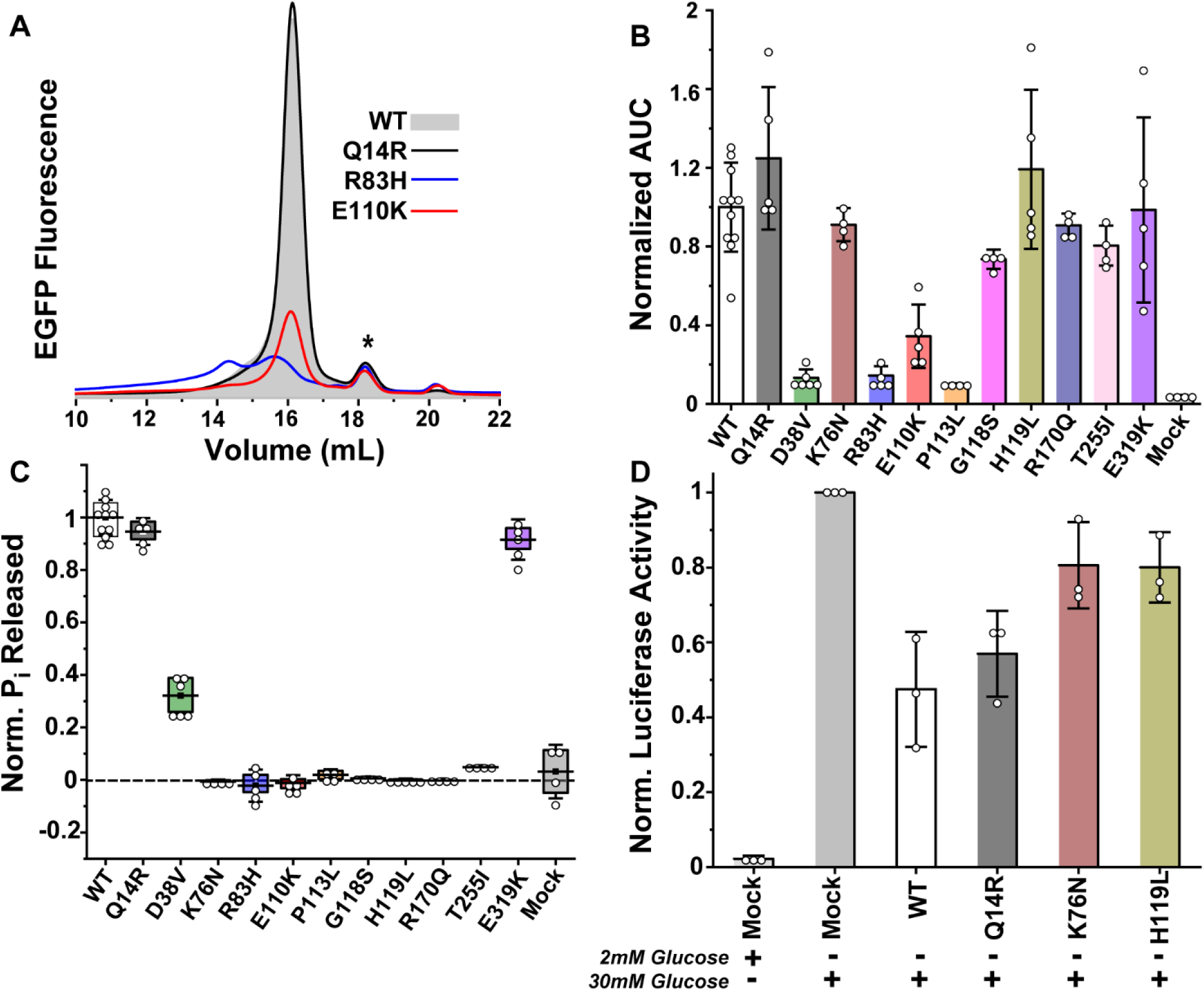
Expression and activity profiles of GSD type1a variants from transfection of adherent cells. (A) Representative FSEC traces illustrate the range of chromatographic behaviors, including reduced expression (E110K) or the presence of misfolded species (R83H). The (*) identifies a minor population of cleaved EGFP. (B) Expression levels, quantified by the area under the curve in FSEC traces, is dependent on the variant. The bar graph captures the average ±standard deviation of multiple independent experiments. (C) The corresponding variant catalytic capacity normalized to its enzyme concentration obtained from (B) is plotted relative to the WT. (D) *In situ* assay of G6PC1 variant activity stimulated by glucose in adherent 832/13 rat islet-derived cells confirms compromised activity of the K76N and H119L variants relative to functional constructs. The metrics in B-D were analyzed by one-way ANOVA and Tukey tests (SI Appendix, Table S1).

Apart from D38V, *in vitro* hydrolysis measurements of LMNG-solubilized G6PC1 with a saturating concentration of G6P revealed a binary enzyme activity profile (Fig 4c). Not surprisingly, the absence of G6P hydrolysis in R83H and P113L was linked directly to either absent or poor expression of folded protein. Even though E110K expressed better than D38V, catalysis was abolished. Likewise, K76N, G118S, H119L, R170Q and T255I were catalytically inactive despite similar expression as WT. We further confirmed the impaired activity of K76N and H119L with an *in situ* luciferase reporter assay that indirectly measures G6Pase activity within intact cells (56, 57). This assay capitalized on a cell line derived from rat islets, 832/13, which displays limited endogenous G6Pase activity. These cells express a transfected rat *G6PC1-luciferase* gene fusion when robustly induced with glucose, stimulating G6P production via glucokinase activity. In the absence of hydrolysis, G6P binds to the luciferase promoter to drive fusion gene expression (Mock, Fig 4d). As expected, co-transfection of functional WT or Q14R blunted glucose-induced fusion gene expression by mediating G6P hydrolysis and reducing glycolytic flux (Fig 4d). On the other hand, cells co-transfected with the K76N and H119L variants behaved similarly to the mock control, consistent with these variants being catalytically impaired (SI Appendix, Table S1).

### Active site contraction induced by GSD type 1a variants

The particularly striking observation that K76N, G118S, H119L, R170Q and T255I retained WT-like expression properties demonstrated that these substitutions are structurally tolerated, precluding a clear correlation with the predicted ΔΔG. Given that energetic disruptions to G6P binding were unequal among these variants (Fig 3), we considered that the molecular consequences of these substitutions may be conferred to the protein backbone. Accordingly, we observed variant-dependent reduction in the solvent accessible surface area (SASA) of the active site calculated by VMD. This reduction was characterized by changes to the ensemble average and breadth of SASA distributions relative to WT and Q14R (Fig 5a), suggesting contraction of the active site. These changes in SASA correlated with average pairwise residue-residue distances in these variants, which highlighted regions of constriction predominately localized to the large luminal loops between TMs 2/3 and TMs 6/7 that contribute residues to the active site (SI Appendix, Fig S8). The restructuring of the active site could be visualized from the perspective of a putative substrate permeable portal composed mostly of residues from TM1 and the large luminal loops (Fig 5b). Selected to represent the mean SASA for each variant (dashed lines in Fig 5a), the frames capture displacement of active site residues and variable degrees of portal collapse. As a corollary, dynamic sampling of the protein backbone was altered relative to WT over the simulation time course. Relative changes in backbone dynamics reported by root mean square fluctuations (ΔRMSF) not only occurred near the active site, but also extended into the transmembrane core (SI Appendix, Fig S9). Collectively, the computational analysis emphasized the potential for active site variants to drive structural alterations that may be causative or degenerate with other mechanisms of catalytic impairment, such as reduced stability as discussed below.

**Figure 5.**
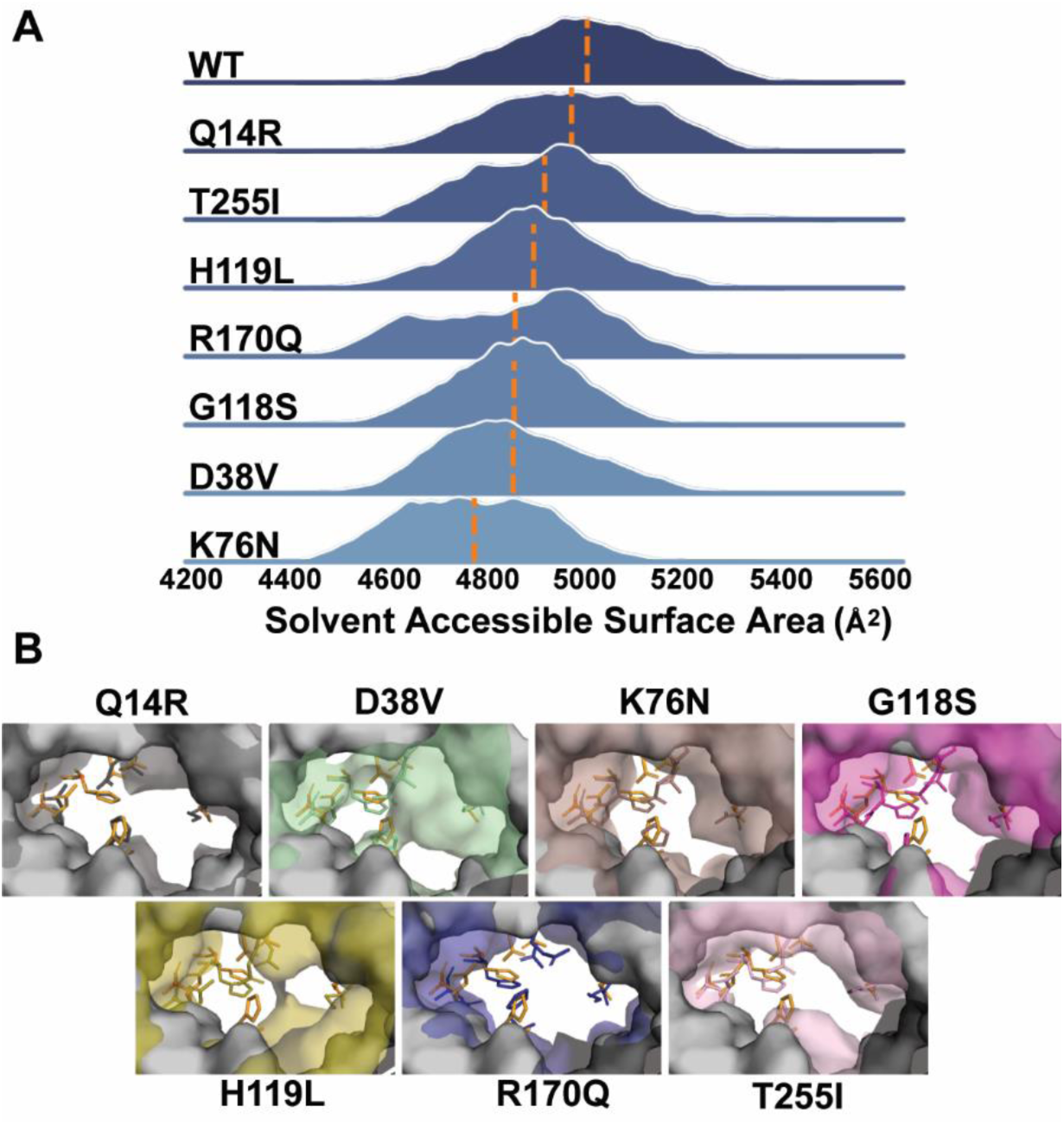
Active site solvent accessible surface area calculated in VMD. (A) The distribution of solvent accessible surface area for each variant relative to WT suggested contraction of the active site. The mean of the distribution is demarcated by a dashed line. (B) View of active site residues through a portal captured side chain repacking and portal collapse at the mean.

### Determinants of reduced expression and hydrolytic capacity of D38V

D38V was the only disease-linked variant that retained catalytic activity significantly above background (Fig 4c). Yet the hydrolysis screen indicated that this variant retained only ∼30% of the Pi release capacity from the equivalent concentration of WT enzyme.Δ The experimental and computational analysis suggested that both expression of folded protein (Fig 4b) and access to the active site (Fig 5) was attenuated strongly. To ascertain more specifically how these observables may reflect structural and catalytic properties, we sought to define turnover kinetics and stability of D38V when purified into LMNG detergent micelles. Unlike Q14R, which displayed nearly identical saturation kinetics as the WT (39), the specific activity of D38V was reduced three-fold without a shift in *K*_M_ (Fig 6a). Thus, the diminished hydrolytic capacity of D38V was a direct consequence of a lower Pi release rate.

**Figure 6.**
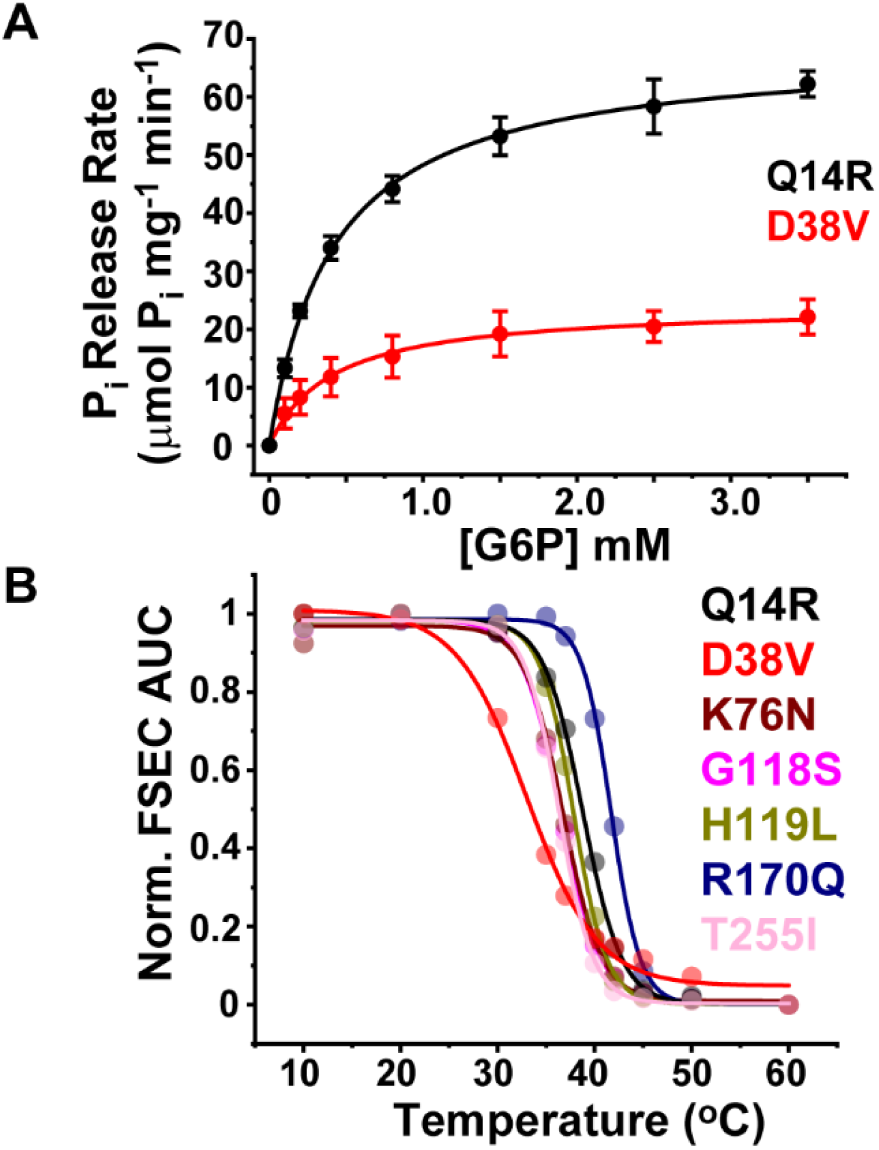
Saturation kinetics of D38V correlates with reduced thermostability. (A) Michaelis-Menten kinetics of G6P hydrolysis indicates that the specific activity of GSD type1a variant D38V (*V*_max_ = 24.5 ±1.9 μmol mg^-1^ min^-1^; *K*_m_ = 0.472 ±0.246 mM) is reduced threefold relative to the WT-like Q14R variant (*V*_max_ = 68.4 ±3.0 μmol mg^-1^ min^-1^; *K*_m_ = 0.413 ±0.017 mM). Solid lines are fits of the data using a single binding site model. (B). The left shift in the D38V melting curve indicates reduced thermostability of D38V relative to other well-expressed variants. Solid lines are fits of the data assuming a simple dose-response model to determine T_M_, which is reported in Table 1.

While a more constricted active site may contribute to the depressed *V*_max_, reduced substrate turnover may also be associated with compromised structural stability. To test the computational prediction implicit of the ΔΔG calculations (Table 1), we measured the thermostability of D38V and compared it to that obtained for other variants. Thermostability was determined by a chromatography-based approach wherein purified protein samples subjected to a thermocycler heating protocol were analyzed for soluble protein content by FSEC monitoring intrinsic Trp fluorescence (39, 58). Plotting the elution peak integral as a function of temperature produced a curve characterized by a melting temperature, T_M_. As shown in Figure 6b, the D38V melting curve was strongly left-shifted relative to Q14R and the other well-expressed disease-linked variants, which was indicative of a loss of heat tolerance. Non-linear least squares fitting of the data confirmed a > 3 °C T_M_ differential between D38V and all other variants (Table 1). Interpreted as a reporter of ΔG*_unfolding_*, this phenomenological analysis of the D38V variant was consistent with compromised thermodynamic stability. Thus, the reduced thermostability of D38V not only rationalized the lower abundance of folded G6PC1, but also correlated with observed catalytic impairment.

## Discussion

Systematic exploration of protein structure and function is widely recognized as essential to defining pathogenic mechanisms arising from missense mutations and that reliable protein models are a prerequisite for contextualizing such biochemical data (59–61). Homology models derived from high resolution structures of related proteins have been proposed to serve as sufficient templates for the interpretation of disease-causing variants (62, 63), although AF2 models are expected to further improve the accuracy of the predictions (64). Conflicting reports of AF2’s sensitivity to missense mutations (65–67) have casted doubt on uncovering either strong correlations or predictive outcomes between WT and variant sequences for a given structural metric or phenotype. However, a recent evaluation of AF2 applications showed that high accuracy AF2 models (pLDDT > 90) matched or improved the correlation between experiment-derived ΔΔG and ΔΔG from structure-based predictors, such as that available with Rosetta, relative to a structure obtained from experiment (64). We show here that combining a high accuracy AF2 model of G6PC1 with orthogonal computational and experimental approaches supports mechanistic insight of critical catalytic interactions and characterization of molecular disruptions imposed by known disease-linked missense mutations.

We find that the AF2 G6PC1 model is highly stable in the simulated *in vivo* environment and that the two stereoisomers of G6P make favorable electrostatic and van der Waals interactions with a network of side chains in the putative active site. This active site, outlined by the consensus phosphatidic acid phosphatase sequence motif, demonstrates similar packing of conserved side chains observed in crystal structures of evolutionary-distant PAP2 superfamily members with bound quaternary ions. Despite subtle repacking of active site side chains in the MD simulations to optimize G6P interactions, the overall conformation of G6PC1 remains intact, suggesting that the AF2 model represents a relevant intermediate in the catalytic cycle. In particular, the tertiary organization of the active site is well suited to mediate hydrolysis chemistry in accordance with the presumed model mechanism (38). While R83 and R170 are arranged to stabilize the substrate and transition state via hydrogen bonds, both H119 and H176 are primed to function as the proton donor and phosphate acceptor, respectively (68).

Numerous phosphorylated substrates for G6PC1 have been identified (7), so the discovery that binding of the β-G6P anomer would be favored by up to three orders of magnitude under equilibrium conditions (*K* ≍ 3300 at 37 °C) was unexpected. Since spontaneous mutarotation of G6P can occur in solution, selectivity for the β-anomer was thought to be exclusively conferred by the cognate SLC37A4 transporter (69). We speculate that preference for β-G6P is associated with sugar pucker angle sampling providing favorable interactions with the G6PC1 active site regardless of the ring conformation (SI Appendix, Fig S1e).

The *in-silico* mutagenesis, facilitating both side chain interaction energy analysis and ΔΔG predictions, largely confirms the anticipated contribution of conserved active-site residues in substrate coordination, but also suggest multiple modes of inhibition that are not necessarily mutually exclusive. For instance, the charge reversal variant E110K substantially increases the binding energy with G6P, which indicates that this variant traps the substrate within the positively charged active site. Moreover, this observation also implies that the native Glu participates in Pi release. This could be achieved mechanistically through dynamics of salt bridge formation and rupture between E110, R170, and R83 over the course of a catalytic cycle, perhaps operating as a modulator of active site polarity to trigger product release. In turn, the R170Q mutation would be expected to compromise salt bridge formation (SI Appendix, Fig S5) in addition to lost electrostatic interactions with the substrate (Fig 3).

Interestingly, the protein stability predictors suggest divergent effects of E110K and R170Q on ΔG*_unfolding_* (Table 1). In concert with the predictions, the destabilizing E110K variant displays highly attenuated expression of folded protein whereas the stabilizing R170Q (supported by experiment, Table 1) reports similar expression as WT. For most variants, a general pattern emerged in which predicted destabilizing substitutions caused either reduced expression or misfolding while neutral or stabilizing substitutions retained WT-like expression profiles. However, this pattern was not explicitly predictive of expression levels with respect to the value of ΔΔG, nor could ΔΔG be assigned as a predictor for activity. For example, while K76N is expected to be moderately destabilizing (+1.3-2.8 kcal/mol), the variant is non-functional likely due to impaired G6P binding (Fig 3) combined with induced structural distortions (Fig 5) despite WT-like expression levels. On the other hand, D38V is predicted to be similarly destabilizing (+2.5-3.7 kcal/mol), and expression is reduced markedly, but it retains a fractional yet significant level of activity relative to WT.

The collective body of work highlights the complexity for which missense mutations found in the active site of G6PC1 conspire to sabotage function via multiple mechanisms. In some cases, the primary means of mechanistic disruption is straightforward, such as for R83H and P113L that induce thermodynamic instability manifested as gross misfolding. However, a confluence of factors mediates impaired catalysis for most active site variants examined here, requiring integration of computational approaches with robust experimental protocols to develop mechanistic models. While thermodynamic instability certainly contributes to expression patterns, altered energy barriers to G6P binding or access to functional intermediates along the reaction coordinate (ΔG°) generate catalytically trapped states, which may be reflected in changes to backbone fluctuations or induced structural distortions. In contrast, Q14R and E319K, variants of uncertain clinical significance, were found to have a benign phenotype in this study and so would be expected to not confer disease. Since the variants explored here are but a small subset of the 72 known GSD type 1a missense mutations in G6PC1, this study supports further characterization of the full complement of variants using the AF2 structural model as a tractable template.

## Materials and Methods

### Modeling of Apo G6PC1 in a simulated ER membrane bilayer

The structural model of mouse G6PC1 (Uniprot P35576) was acquired from the publicly available AF2 database (https://alphafold.ebi.ac.uk/). The initial equilibrium simulation was constructed using the CHARMM-GUI web server (70). An approximate ER lipid bilayer composition was chosen consisting of 55% POPC, 20% POPE, 10% POPI, 5% POPS, 5% cholesterol, and 5% sphingomyelin for both membrane leaflets. The position of G6PC1 relative to the bilayer was computed using the Orientation of Proteins in Membranes server (OPM) being facilitated by CHARMM-GUI to ensure proper placement (71, 72). The pKa of titratable residues was calculated by PropKa3.1. H119 was protonated (HSP) as predicted by its catalytic role as a proton donor in the reaction mechanism. All other histidine residues were assigned neutral (HSD).

The model was relaxed from its initial state over a series of equilibration steps with decreasing restraints using NAMD2 and the CHARMM36 force field (53, 73). First a 10,000 step minimization was performed after which a slow heating protocol of increasing the system temperature by 25 K every 10 ps until 310 K was reached. During heating and the first 2.5 ns of equilibration there was a 1 kcal/mol/Å^2^ harmonic restraint was placed on lipid head group and protein backbone atoms to allow the lipid tails to melt. This was followed by 2.5 ns of simulation with a 1 kcal/mol/Å^2^ restraint only on the protein backbone, and a final 2.5 ns of simulation with a 0.5 kcal/mol/Å^2^ restraint on protein alpha carbons. Five nanoseconds of unrestrained equilibration was then performed to allow for switching to NPT and subsequent relaxation of the system.

Simulation replicas were generated after equilibration using the Membrane Mixer tool in VMD in order to sufficiently sample the initial lipid configuration of the system (74, 75). Production MD runs were carried out using NAMD3 with a timestep of 2 fs and a long-range interaction cutoff of 14 Å (54). In production runs, each replica was simulated for 100 ns resulting in a cumulative total of 300 ns of trajectory data.

### *In silico* mutagenesis and thermodynamic predictions

ΔΔG calculations were performed to predict the thermodynamic effects of missense variants using Rosetta following the method described in Ref (47). Briefly, the model was oriented in an implicit membrane environment by aligning it to the OmpLA structure provided in $ROSETTA/main/demos/public/mp_ddg/inputs/1qd6_tr_C.pdb using UCSF Chimera 1.16 (76). The spanfile was generated using the consensus server at TOPCONS.net (77) and edited to optimally specify the membrane-spanning helices. The structure was relaxed into Rosetta’s franklin2019 score function using 1000 independent runs of the rosetta_scripts executable version 2020.37.61417 with the membrane_relax.xml protocol. The lowest-scoring relaxed decoy was then used as the input for the predict_ddG.py script with a repack radius of 8.0 Å. The predict_ddG.py script was adjusted to run under pyrosetta-2021.49 release r305 (Python 3.8.12, Conda version 5.0.1) inside a Slurm array of 357 jobs on Vanderbilt’s ACCRE cluster to assess all 20 possible mutations at each position in the protein. The ΔΔG values of disease-associated variants were compiled from the output files using GNU Awk 4.0.2, mapped onto the structural model using Pymol 2.5.4 (78) (SI Appendix, Fig S3) and reported in Table 1. The error associated with these predictions is expected to be within 1 kcal/mol on average (48, 50, 79). Therefore, any value greater than 2 kcal/mol was considered destabilizing, whereas any value less than −2 kcal/mol was considered stabilizing.

### Preliminary docking of G6P

Crystal structures of bacterial PAP2 type lipid phosphatases (PDB 6EBU, 6FMX and 5JKI) and vanadium containing chloroperoxidase (PDB 1IDQ) were aligned with the AF2 G6PC1 model using the catalytic Arg, His and Lys side chains from the signature phosphatase sequence motif. In MATLAB (MathWorks), the 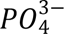 moiety of the α- and β-G6P anomers (also obtained from crystal structures) were transformed onto the atomic coordinates of the quaternary ions of 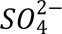, 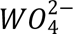, and 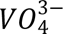 from the aligned structures yielding 95 potential configurations of G6P. Any G6P model within 1.2 Å of the AF2 model, whether in side chains or backbone atoms, was discarded to prevent van der Waals clashes. From the remaining G6P models, two were chosen as representative α- and β-G6P for further MD simulations.

### Equilibrium simulations with docked G6P

A similar equilibration protocol to the Apo simulations was followed with the notable exception of including G6P restraints with the same force constant as the protein heavy atoms throughout restrained equilibration. Parameters for G6P were obtained from phosphorylated sugar parameters in the CHARMM36 force field. The same initial lipid configuration as in the Apo replicas was used to seed the docked simulations and each was simulated for 250 ns of production MD for each G6P anomer.

From the initial WT state, nine mutants were also generated (Table 1) using the Mutagenesis tool in VMD. In each mutant state, the system charge was adjusted accordingly by reionization to ensure a net charge of 0. Each replica was equilibrated as above for the WT systems and simulated for 250 ns resulting in a cumulative total of 15 µs of trajectory data.

All computational analysis was performed using in-house TCL scripts, the MDAnalysis python package or tools that are built into VMD. This includes solvent accessible surface area calculations via the SASA tool (74, 80).

### Free Energy Perturbation calculations

Free energy perturbation (FEP) was performed using a representative model for both G6P anomers. In each system the substrate molecule was duplicated and placed over 25 Å away from the protein in the bulk solvent. The resulting system was re-minimized and then gently equilibrated for 30 ns to ensure accurate results. The protein backbone was restrained as well as the G6P phosphorous atom throughout the simulation with a force constant of 1 kcal/mol/Å^2^ to allow for side chain rearrangement and substrate internal sampling to still occur. Each FEP protocol consisted of 50 windows of lambda values ranging from 0 to 1 and incrementing/decrementing by 0.02 each window. Each window was simulated for 1 ns for a total of 50 ns per FEP simulation set in each direction.

Free energies were extracted from the trajectories using the Analyze FEP plugin in VMD. Gram-Charlier expansion was set to 0 and the Bennet Acceptance Ratio (BAR) estimator was used to compute error. Forward and backward paths were plotted to ensure convergence of the simulation set. Relative binding free energy was then calculated and reported.

### Screening G6PC1 expression and activity *in vitro*

Measurements of expression and G6P hydrolysis following transfection of adherent HEK293SG cells (*N*- acetylglucosaminyl-transferase I-negative; ATCC CRL-3022) followed the previously published methods (39). Briefly, WT mouse DNA (accession number NM_008061) was cloned into the pJPA5 MOD expression vector. EGFP was fused to the C-terminus of G6PC1 with a linker that included a thrombin protease recognition sequence. Mutations were introduced via site directed mutagenesis using complementary oligonucleotide primers, and DNA sequencing confirmed the presence of the desired mutation and the absence of unwanted changes. Adherent semiconfluent HEK293SG cells cultured in DMEM:F12 medium supplemented with 10% FBS were transfected in 6-well plates with 2 μg/well of plasmid DNA complexed with Lipofectamine 2000 reagent (Invitrogen). The cells were incubated for 48 hrs at 37 °C under 7% CO_2_. Expression was confirmed by visualizing cell epifluorescence on a Zeiss Axio Zoom.V16 fluorescence stereo microscope. Epifluorescence intensity was quantified for mock (H_2_O), WT and variants across multiple independent transfection experiments (n=3-7) using the ImageJ application (81) running the StarDist plugin.

Following harvest, the cells were solubilized in 300 μL buffer containing 50 mM Tris-HCl pH 8.0, 150 mM NaCl, 1 mM EDTA, 5 mM (0.5% w/v) LMNG and 5mM PMSF. Insoluble material was removed by ultracentrifugation at 105,000 rcf for 20 mins. Supernatant containing solubilized G6PC1-EGFP was injected onto a Superose6 Increase 10/300 GL column equilibrated in 50 mM Tris-HCl pH 8.0, 150 mM NaCl, 1 mM EDTA and 0.01% (w/v) LMNG buffer. The column was attached to an Agilent 1260 Infinity II chromatography system equipped with a fluorescence detector and a temperature-controlled multi-sampler. Sample elution profiles monitored EGFP fluorescence (Ex 475 nm, Em 515 nm). The enzyme concentration was estimated by integration of the elution peak obtained from 15-17 mL and normalized to the WT for each transfection experiment (n=4-9). Variability in the calculated expression level (Fig 4) was a function of transfection efficiency compounded with detergent solubilization efficiency for each experimental iteration. One-way ANOVA and the Tukey test were used to assess statistical variation in the mean epifluorescence and FSEC peak area datasets at the 0.05 level (SI Appendix, Table S1).

Phosphohydrolase activity was measured by diluting 20 μL of LMNG-solubilized enzyme into a final volume of 150 μL 50 mM Tris/Mes pH 6.5, 50 mM NaCl, 0.2mM LMNG in the presence or absence of 1.5 mM G6P and mixed on ice. The reaction was transferred to a 30 °C water bath for five minutes to stimulate G6P hydrolysis. The reaction was quenched with 150 μL of 12% (w/v) SDS and vortexed. The amount of Pi released was determined from a colorimetric assay relative to a Pi standard curve as previously described. Absolute Pi released was normalized to enzyme concentration determined from the FSEC peak area and then scaled relative to the WT. Reactions were performed in triplicate per biological repeat (n=4-9), and the means subjected to ANOVA and Tukey tests (SI Appendix, Table S1).

### Measurement of glucose-6-phosphatase (G6Pase) activity *in situ*

G6Pase activity was measured *in situ* as previously described (56). Briefly, semi-confluent 832/13 cells in 3.5 cm diameter dishes were co-transfected with 2 μg of a *G6pc1*-firefly *luciferase* fusion gene construct, 0.5 μg of SV40-*Renilla luciferase* (Promega, Madison, WI) and 1 μg of an expression vector encoding WT or mutated G6PC1-EGFP, using the lipofectamine reagent (InVitrogen, Waltham, MA). Following transfection, cells were incubated for 18-20 hours in serum-free medium supplemented with 2 or 30 mM glucose. Cells were then harvested using passive lysis buffer (Promega, Madison, WI) and both firefly and *Renilla* luciferase activity were assayed using the Dual Luciferase Assay kit (Promega, Madison, WI). To correct for variations in transfection efficiency, the results were calculated as a ratio of firefly activity to protein concentration in the cell lysate. The four G6PC1 constructs (WT, Q14R, K76N and H119L) were measured in duplicate with n=3 independent biological replicates. The means were subjected to ANOVA and Tukey tests (SI Appendix, Table S1).

### Expression and purification of G6PC1 from Sf9 insect cells

Heterologous expression and purification of Q14R and D38V followed the previously published protocols with minor adjustments (39). Briefly, G6PC1 in the pFastBac1 vector was expressed as an EGFP fusion that contains a C-terminal His_8_ tag via baculovirus transduction of Sf9 insect cells for 72 hrs at 27 °C. Isolated membranes were solubilized with 5 mM (0.5% w/v) LMNG in 50 mM Tris pH 8, 100 mM NaCl, 10% (v/v) glycerol buffer for 1 hr on ice. Insoluble material was removed by ultracentrifugation at 185,000 rcf for 50 min. The supernatant was filtered through a 0.45 μm syringe filter and mixed with Ni^2+^-NTA Superflow (Qiagen) equilibrated in the same buffer supplemented with 25 mM imidazole for 4 hrs at 4 °C with gentle shaking. The resin was washed with 10 column volumes of buffer containing 100 mM imidazole followed by elution of G6PC1-EGFP with 300 mM imidazole buffer. EGFP was cleaved via thrombin digestion (0.04 NIH units/μg of fusion protein) at 15 °C for 16 hrs. EGFP and thrombin was removed from G6PC1 by gel filtration through a Superose6 Increase 10/300 GL column equilibrated with 50 mM Tris pH 7.5, 150 mM NaCl, 0.2 mM LMNG. Enzyme concentration was determined from absorbance at 280 nm assuming a 90,340 M^-1^ cm^-1^ molar extinction coefficient calculated from the protein sequence. Sample purity was assessed by SDS-PAGE using a 13% acrylamide gel.

### Measurement of G6P hydrolysis and thermostability with purified G6PC1 variants

The Pi release rate was determined from titration of 0.1 μg (2.4 pmol) G6PC1 with G6P at 30 °C for 1 min at pH 6.5 as previously described (39). A linear [G6P]-dependent curve collected on ice was subtracted from reaction carried out at 30 °C. Catalytic parameters were determined from fits of the saturation curve assuming a Michaelis-Menten model in the program Origin (OriginLab). The reported mean values and standard deviations of *K*_M_ and *V*_max_ were derived from analysis of duplicate hydrolysis curves acquired from two independent enzyme preparations for each variant.

The thermostability of each variant was described by the temperature-dependent, irreversible depletion of soluble enzyme as observed by FSEC (39, 58). Purified Q14R or D38V (0.1 mg/mL) was transferred to standard PCR tubes and incubated in a thermocycler at defined temperatures for 10 min intervals. Precipitated material was removed by ultracentrifugation at 105,000 rcf and the supernatant injected on a Superose6 Increase 10/300 GL column equilibrated in 50 mM Tris pH 7.5, 150 mM NaCl, 0.2 mM LMNG buffer. Enzyme elution was monitored by fluorescence (Ex 280 nm, Em 320 nm). Peak area, representing the remaining soluble population, was plotted as a function of increasing temperature and the data fit with a dose-response curve in the program Origin to quantify the melting temperature (T_M_).

## Acknowledgments

This work was supported by the National Institutes of Health grants P41-GM104601 and R24-GM145965 (M.S. and E.T.) and R01-DK132259 (R.O’B.), the Vanderbilt Molecular Endocrinology Training program 5T32-DK07563 (E.M.H.), the American Heart Association 23PRE1017904 (E.M.H.), the Juvenile Diabetes Research Foundation 1-INO-2022-1121-A-N and Vanderbilt University’s Program in the Molecular Basis of Genetic Diseases. We also acknowledge computing resources provided by eXtreme Science and Engineering Discovery Environment (XSEDE) (grant MCA06N060 to E.T.). The authors wish to thank Mr. Michael Mohan for assistance with baculovirus production and protein expression and Dr. Chris Newgard for providing the 832/13 cell line. We thank Dr. Hassane Mchaourab and Dr. Benjamin Brown for a critical reading of the manuscript.

## SI Appendix

**Table S1.** Means testing for G6PC1 variants for biological metrics relative to WT. The mean difference, standard error of the mean and the associated P-value are reported Means that are significantly different at the 0.05 level are colored red

| Variant | Epifluorescence |  |  | FSEC AUC |  |  | *In Vitro* Activity |  |  | *In Situ* Activity |  |  |
| --- | --- | --- | --- | --- | --- | --- | --- | --- | --- | --- | --- | --- |
|  | Mean Diff | SEM | P | Mean Diff | SEM | P | Mean Diff | SEM | P | Mean Diff | SEM | P |
| Q14R | 0.172 | 0.089 | 0.765 | 0.248 | 0.123 | 0.718 | -0.054 | 0.029 | 0.823 | 0.095 | 0.081 | 0.842 |
| D38V | -0.678 | 0.080 | 4.86E-8 | -0.867 | 0.116 | 1.2E-7 | -0.679 | 0.027 | 0 | N.D. |  |  |
| K76N | 0.045 | 0.080 | 0.999 | -0.089 | 0.133 | 0.999 | -1.007 | 0.032 | 1.13E-6 | 0.331 | 0.081 | 0.014 |
| R83H | -0.760 | 0.089 | 4.36E-8 | -0.856 | 0.123 | 4.82E-7 | -1.022 | 0.029 | 7.55E-8 | N.D. |  |  |
| E110K | -0.453 | 0.089 | 6.18E-4 | -0.656 | 0.123 | 1.47E-4 | -1.012 | 0.029 | 7.31E-8 | N.D. |  |  |
| P113L | -0.536 | 0.080 | 4.2E-6 | -0.901 | 0.133 | 9.16E-7 | -0.981 | 0.032 | 0 | N.D. |  |  |
| G118S | -0.168 | 0.080 | 0.671 | -0.265 | 0.133 | 0.738 | -0.994 | 0.032 | 0 | N.D. |  |  |
| H119L | -0.086 | 0.089 | 0.999 | 0.192 | 0.123 | 0.934 | -1.002 | 0.029 | 7.08E-8 | 0.325 | 0.081 | 0.016 |
| R170Q | 0.143 | 0.080 | 0.842 | -0.092 | 0.133 | 0.999 | -1.002 | 0.032 | 1.12E-6 | N.D. |  |  |
| T255I | -0.265 | 0.080 | 0.086 | -0.196 | 0.133 | 0.957 | -0.952 | 0.032 | 0 | N.D. |  |  |
| E319K | -0.165 | 0.089 | 0.811 | -0.014 | 0.123 | 1.000 | -0.085 | 0.029 | 0.191 | N.D. |  |  |
| Mock | -0.979 | 0.080 | 8.39E-8 | -0.961 | 0.133 | 2.30E-7 | -0.968 | 0.032 | 0 | -0.525* | 0.081* | 3.28E-4* |

(\*) Comparison made with WT and mock controls at 30mM glucose

**Figure S1.**
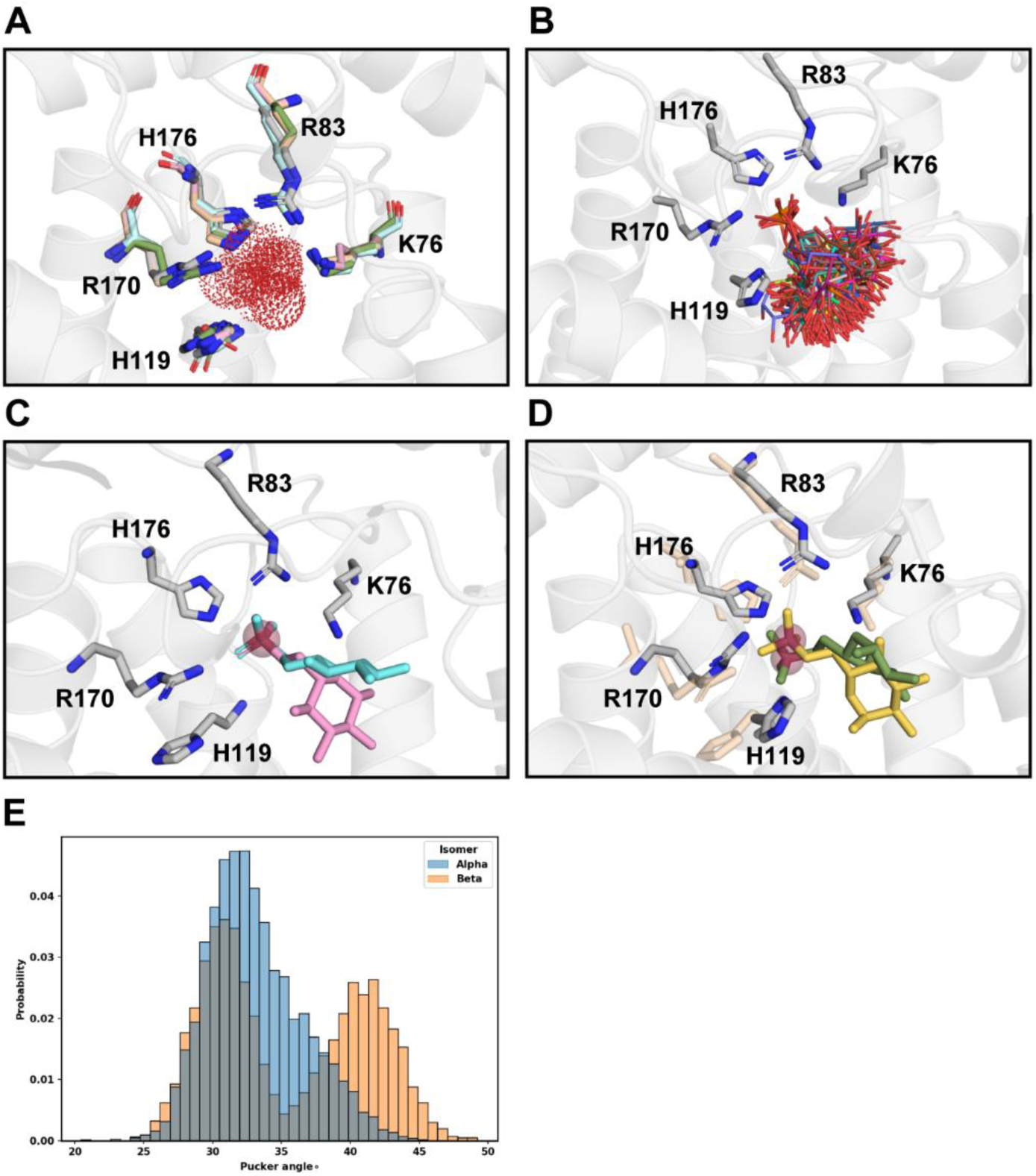
Docking of G6P anomers to the predicted G6PC1 active site. (A) Superposition of the active site from PAP2 crystal structures (PDB ID 6EBU, 6FMX, 5JKI and 1IDQ) with the AF2 G6PC1 model (Uniprot P35576) indicating the conserved side chain projections and atomic positions for surrogate phosphate ions. The residue numbering scheme follows the G6PC1 primary structure. (B) Preliminary docking of 95 G6P structures from both anomers to the consensus position of the quaternary ions in (A). (C) Representative α- (pink) and β- G6P (cyan) models docked to the bilayer-relaxed G6PC1 model. (D) Comparison of G6P anomer positioning and side chain rotamers at t = 0 of the MD simulations. Here, the α- and β-G6P models are yellow and green, respectively. The position of rotamers in the Apo model (tan) are as shown in (C) prior to initiating the equilibrium simulations. (E) Distribution of sugar pucker angles over the course of simulation in WT protein. The bimodal nature of β-G6P correlates with ring flipping observed in simulation.

**Figure S2.**
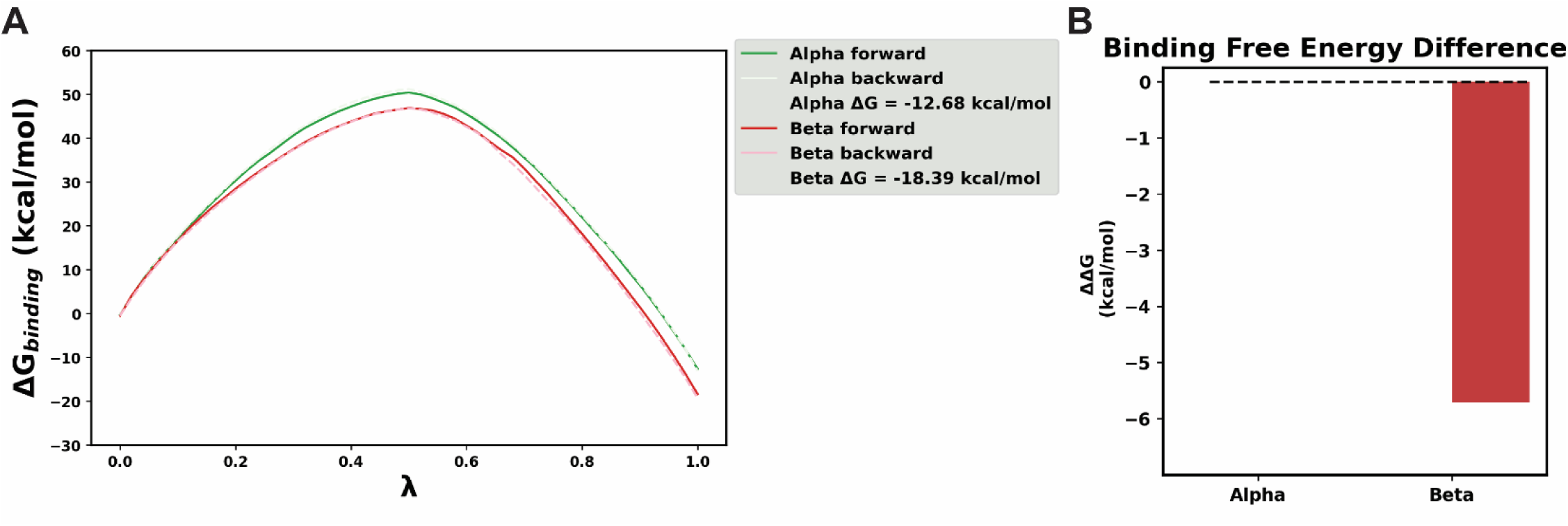
Free energy perturbation calculations of G6P anomer binding. (A) Forward and backward paths for binding of each anomer during a 50-window FEP simulation set. Convergence is observed for both anomers as well as a clean overlay of the forward and backward paths. (B) Relative binding free energy of the anomers.

**Figure S3.**
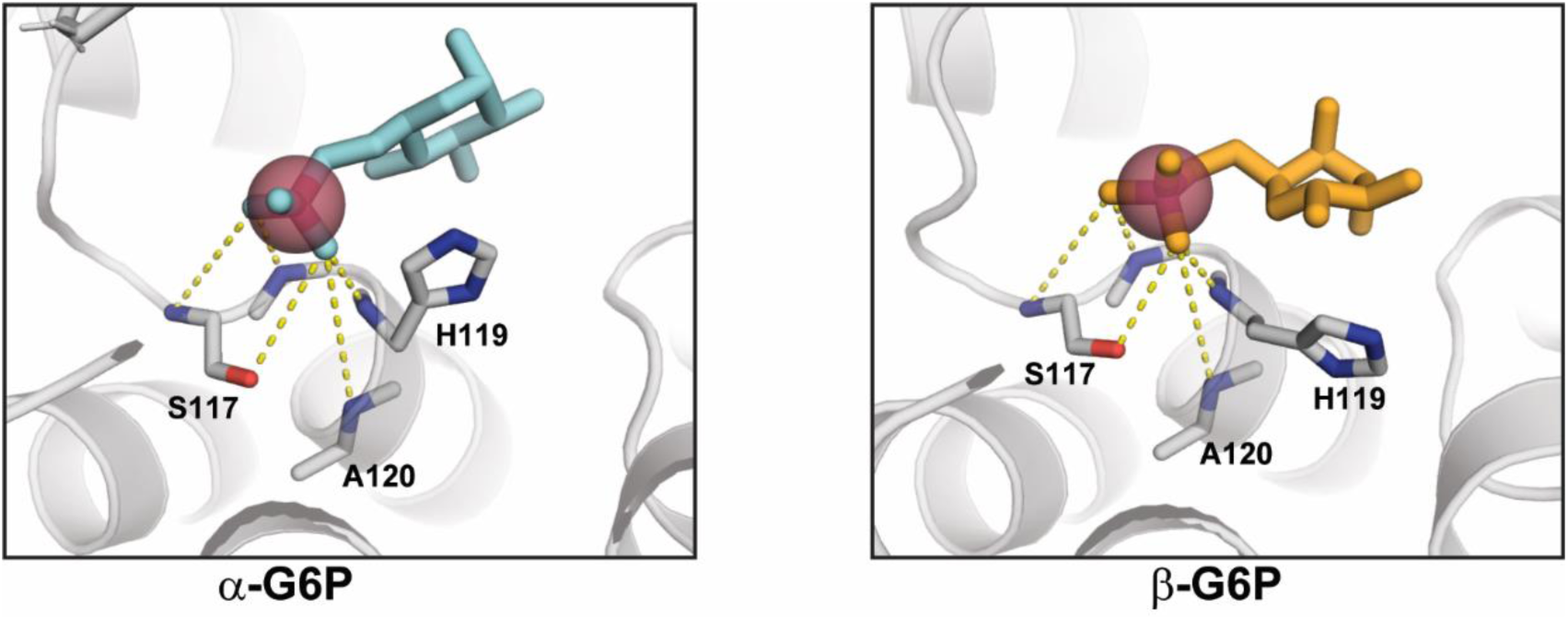
Backbone amides contribute to electrostatic interactions with G6P anomers. Backbone amides of S117, G118, H119 and A120 contribute hydrogen bonding interactions with the phosphate moiety. The frames correspond to those shown in Fig 2c-d.

**Figure S4.**
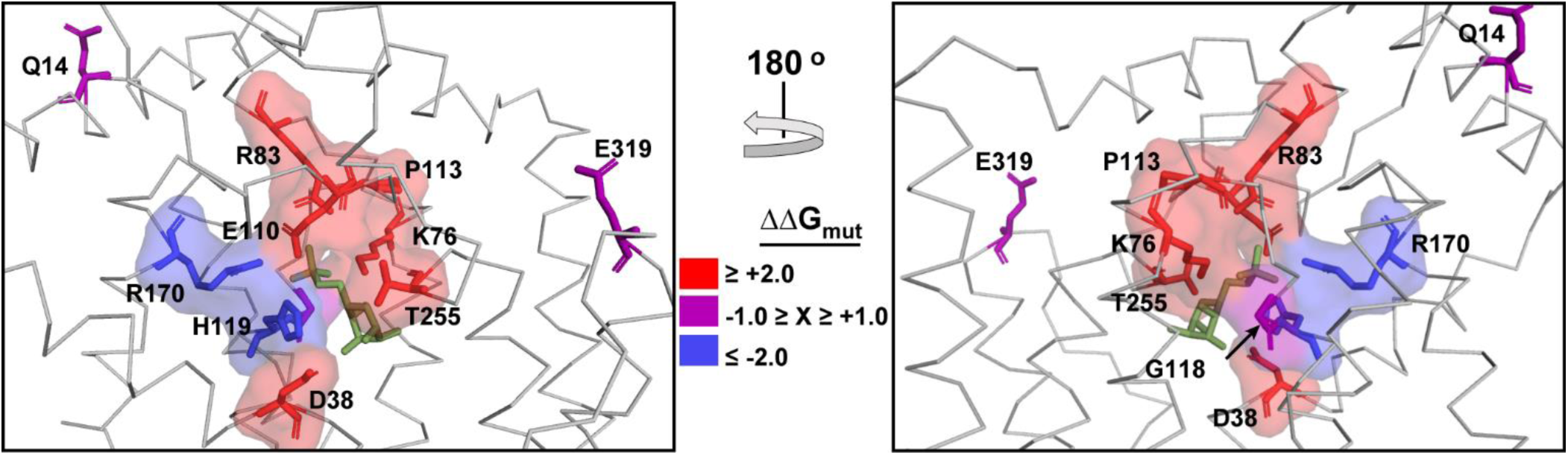
Mapping of Rosetta. ΔΔ**G onto the AF2 G6PC1 model.** Relative location of amino acids associated with GSD type 1a in the active site. Variants predicted to be destabilizing are colored red whereas stabilizing variants are colored blue (Table 1). The position of two variants of unknown significance that reside outside the active site, Q14 and E319, are shown as stick representations.

**Figure S5.**
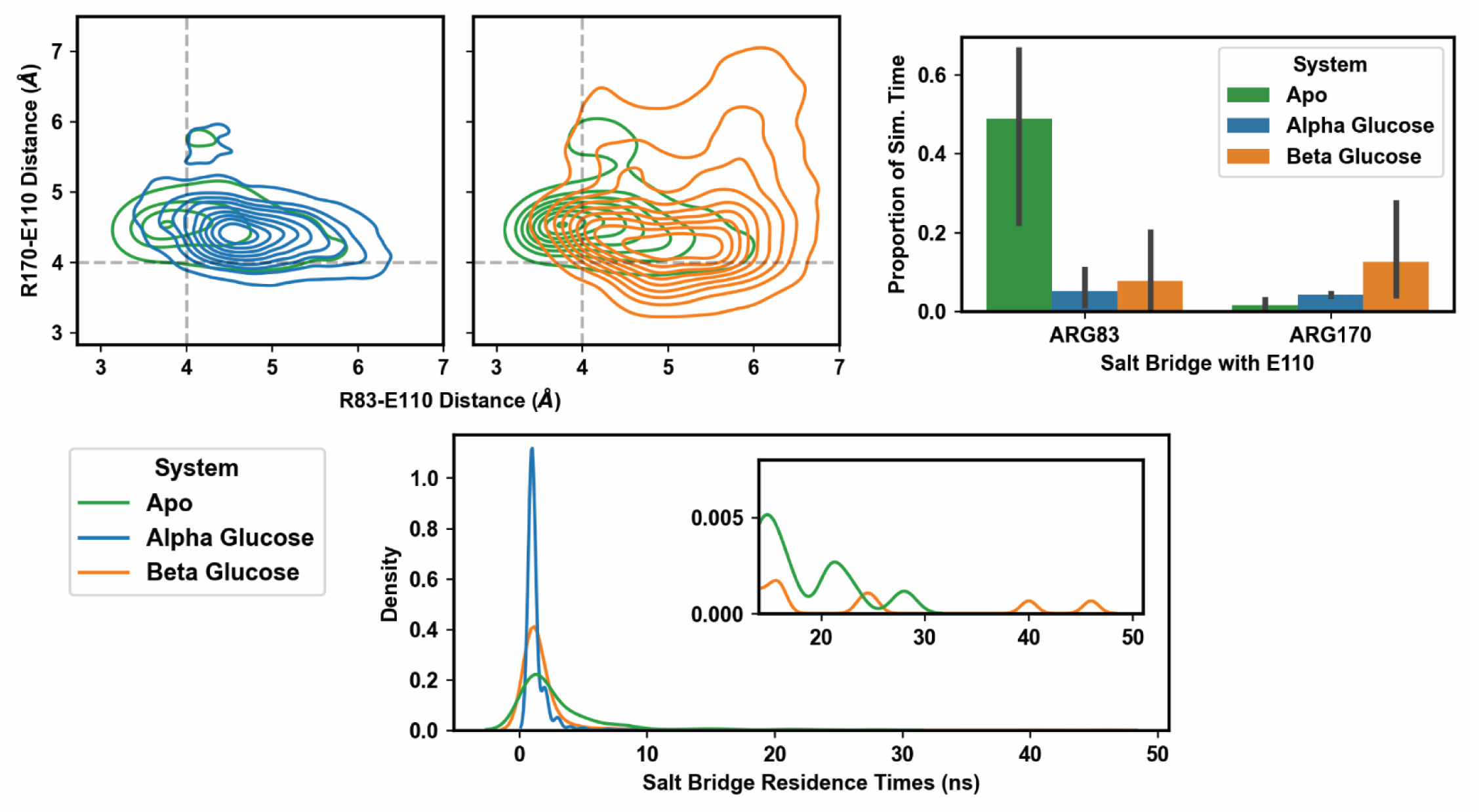
Salt bridge interactions between R170, E110, and R83. The distribution of salt bridge distances in simulations is shown at top left with the R83-E110 and R170-E110 salt bridges mapped on the x- and y-axis, respectively. A 4 Å cutoff for salt bridge distance is displayed as a grey dashed line for each pair. The proportion of simulation time spent in a salt bridge by system is shown at top right. The standard deviation is computed across all three replica simulations for each system and salt bridge. The distribution of salt bridging event residence times by system is shown at bottom. The inset shows rarer, high residence salt bridge events in the Apo and β−anomer−bound systems.

**Figure S6.**
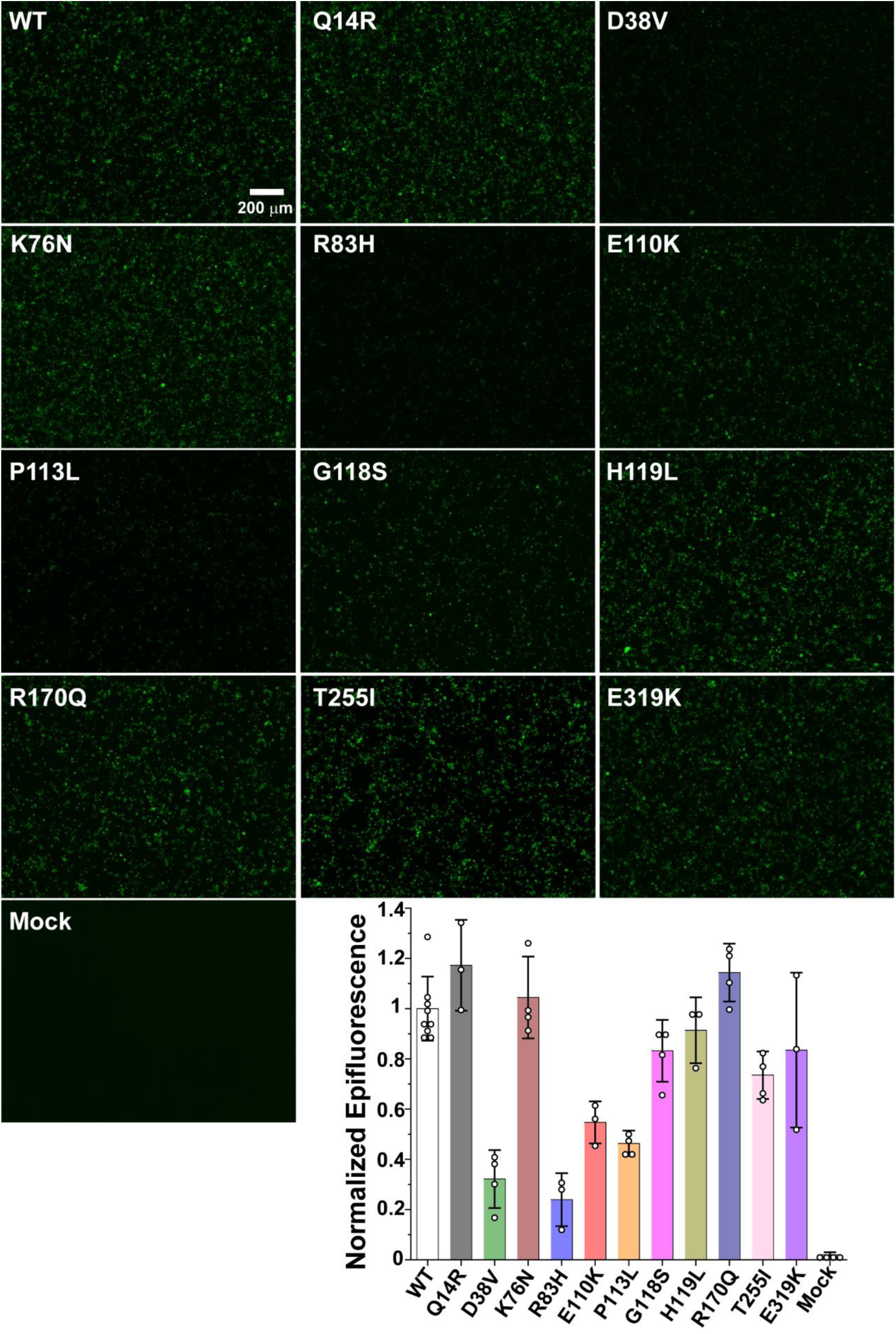
Epifluorescence of adherent HEK293SG cells following transfection of WT and variant G6PC1. Representative images are shown at the indicated scale and the fluorescence quantified in ImageJ software. The pattern of expression is comparable to that observed from FSEC analysis of detergent solubilized enzyme. The quantified fluorescence was analyzed by one-way ANOVA and Tukey tests (SI Appendix, Table S1).

**Figure S7.**
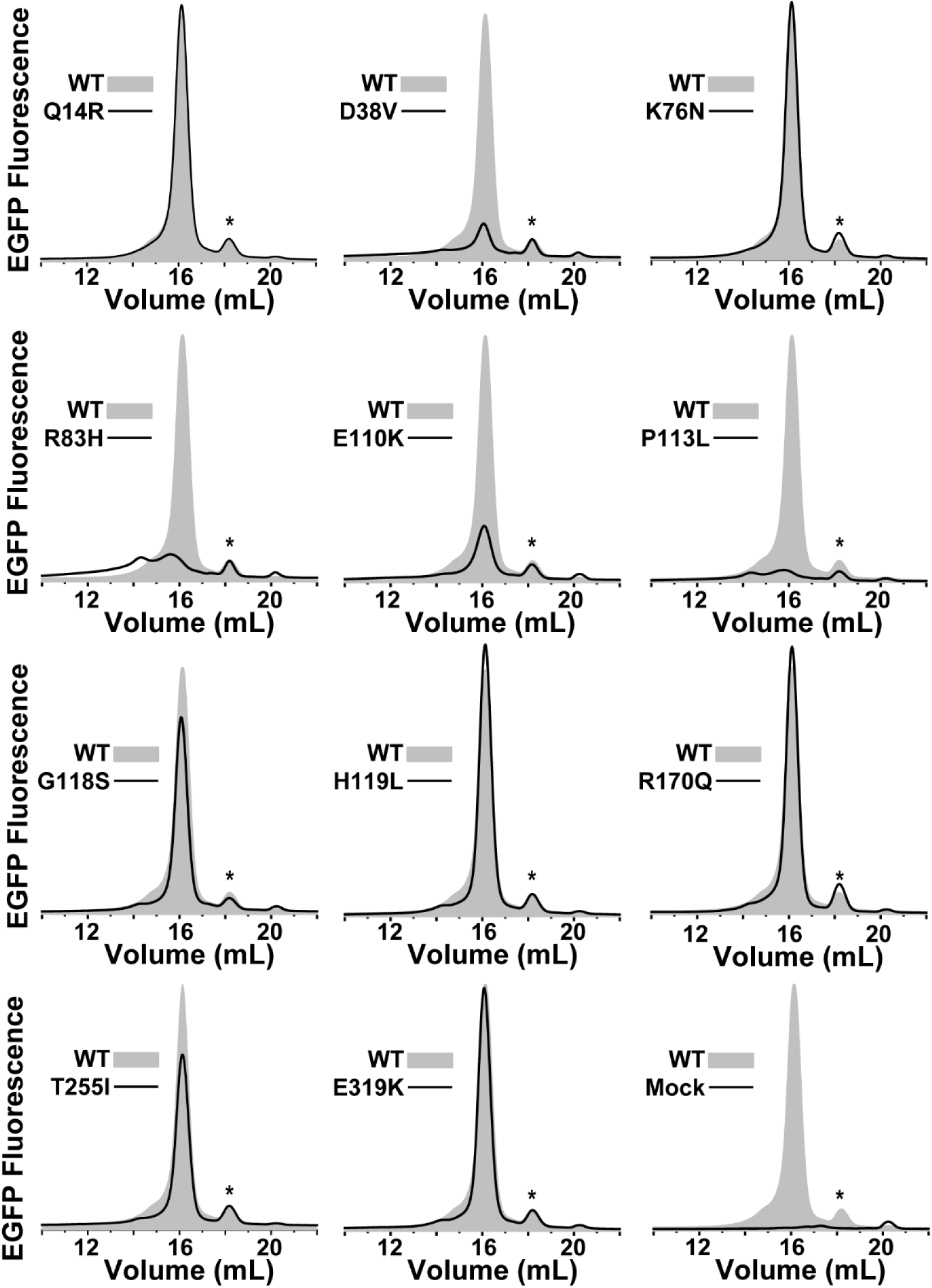
FSEC traces of GSD type1a variants relative to WT. Representative chromatographic profiles of detergent solubilized G6PC1 illustrate the impact of disease linked variants to enzyme expression and folding. The (*) identifies a minor population of cleaved EGFP in the samples.

**Figure S8.**
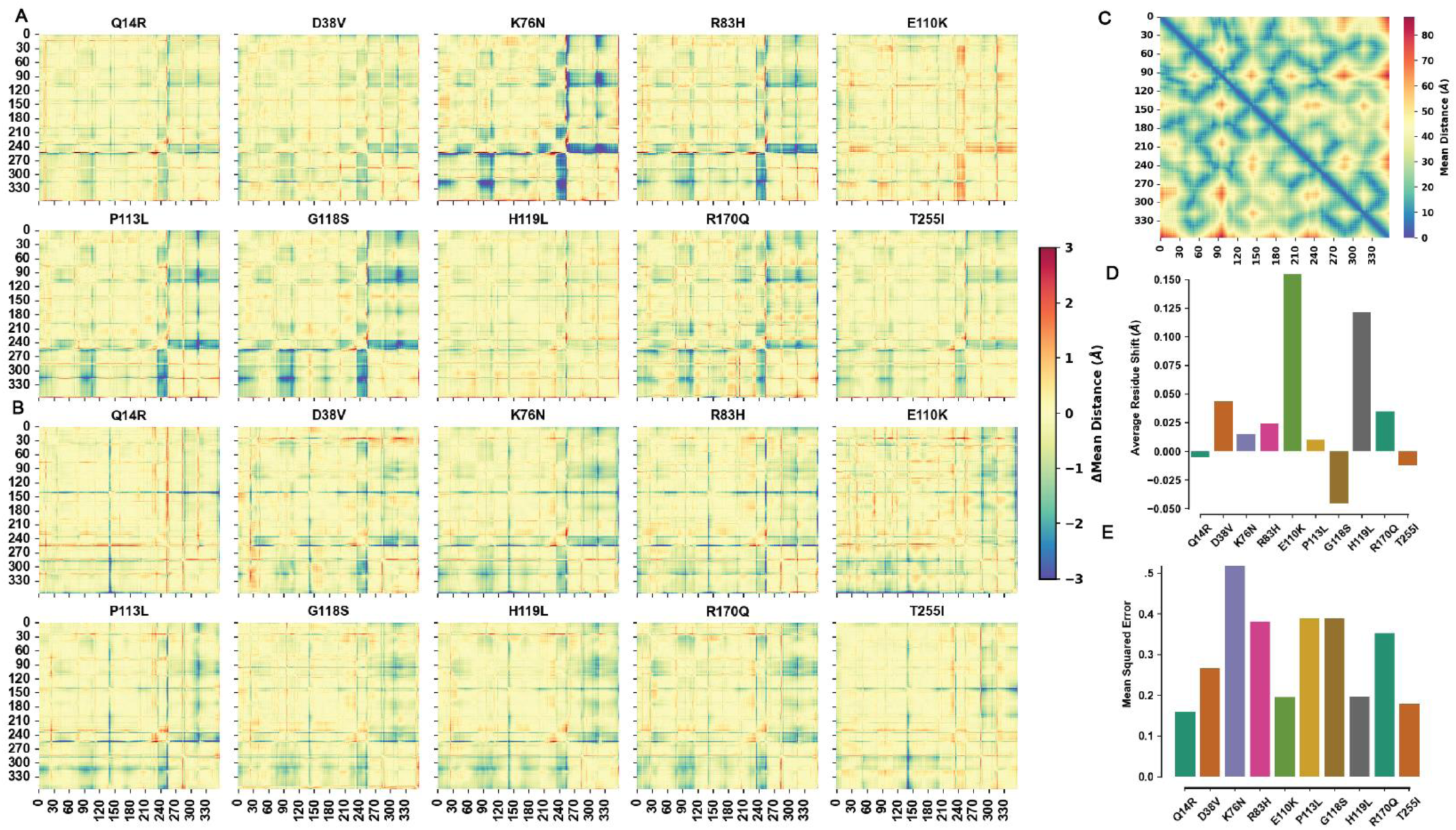
Identification of structural distortions induced by GSD type 1a variants. Δ mean pairwise residue-residue distance matrices for each mutant with bound (A) α-G6P and (B) β-G6P. Matrices were obtained by subtracting the mean pairwise residue-residue distance matrix in WT from values for each mutant. Positive values indicate a given pair of residues are farther apart in the variant, while negative values indicate they get closer together. (C) Mean pairwise residue-residue distance matrix for WT G6PC1. (D) Mean pairwise distance changes over all residues for a given mutant and the associated (E) mean squared error (MSE) identify changes mainly due to backbone dynamics (higher MSE, lower mean distance) or due to induced structural distortions (lower MSE, higher mean distance).

**Figure S9.**
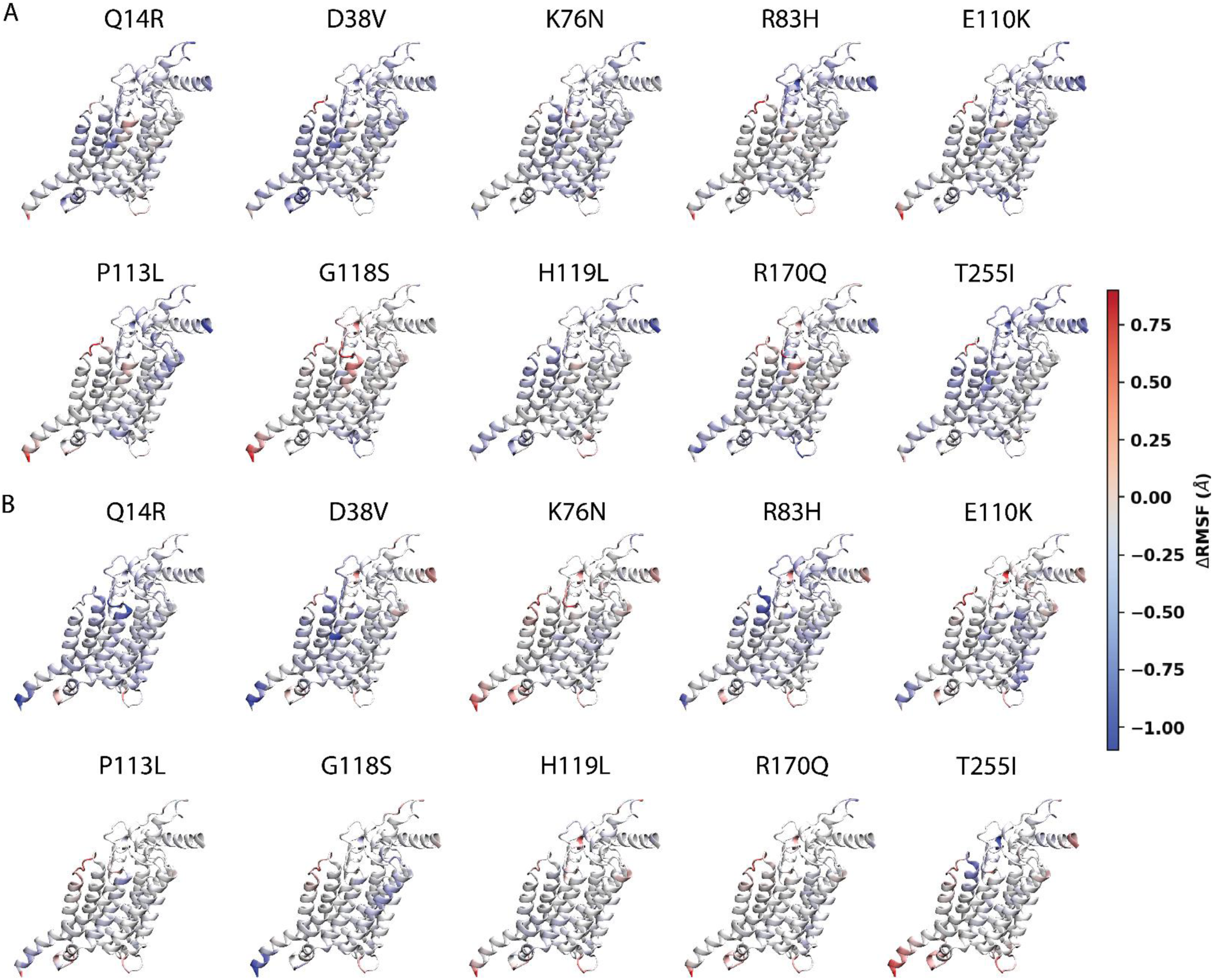
ΔRMSF of GSD type1a variants. Mapping of changes in backbone flexibility induced by each variant and parsed according to G6P anomer. (A) α-G6P and (B) β-G6P. Regions of positive ΔRMSF indicate an increase in flexibility while regions of negative ΔRMSF indicate a decrease in flexibility.

